# Facultative release from developmental constraints through polyphenism promotes adaptively flexible maturation

**DOI:** 10.1101/2021.06.14.448392

**Authors:** Flor T. Rhebergen, Isabel M. Smallegange

## Abstract

The timing of maturation, a critical fitness determinant, is influenced by developmental and energetic constraints, particularly when growth is poor in adverse conditions. Such constraints can be altered through developmental plasticity. Thus, in theory, plasticity in energetically costly sexually selected morphologies can promote life history flexibility in variable environments. We experimentally tested this hypothesis in bulb mites *(Rhizoglyphus robini)* that polyphenically develop as armed fighters with enlarged legs or as scramblers without modified legs. We found that (i) mites enter metamorphosis earlier if they develop as scramblers, (ii) mites accelerate the onset of metamorphosis when they sense resource limitation, and (iii) scrambler expression increases under increased competition for food, enabling males to mature early and escape juvenile mortality. We propose that life history plasticity can evolve through polyphenic release from sexually selected constraints, making the evolutionary dynamics of secondary sexual traits and life history traits, typically studied separately, interdependent.

## Introduction

Age and size at maturity are life history traits with major fitness consequences, and are characteristically plastic^1,2^. This plasticity evolves under environmental variation, as the balance between fitness costs (juvenile mortality risk, generation time length) and fitness benefits (larger body size at reproduction) of postponing maturation shifts with nutritional circumstances. If resources are abundant and growth is rapid, it generally pays to postpone maturation past the stage at which maturation is physiologically possible, to become large. However, if resources are scarce and growth is slow, fitness is optimized by maturing as early as physiologically possible^3–6^. Hence, in favourable circumstances, optimal age and size at maturity are generally determined by ecological opportunities and the benefits of being large^7^. In unfavourable circumstances, optimal age and size at maturity are often more strongly influenced by developmental demands and energetic constraints^3,6,8,9^, which can be altered through developmental plasticity^10^.

In the majority of animal species, maturation occurs through metamorphosis^11–13^, during which several energetically expensive processes compete for resources^14^: organisms restructure their bodies, develop reproductive organs, and sometimes also develop secondary sexual traits to be used as weapons or ornaments in competition for mating opportunities^15,16^. Secondary sexual traits are not only typically costly to produce and maintain^17–20^, but often also developmentally plastic and nutrition-sensitive: poorly-fed individuals often develop proportionally smaller sexual traits, or no sexual traits at all (polyphenism)^21–25^. This suggests that individuals may not always be developmentally obliged to acquire a large resource budget and produce sexual traits. Instead, individuals may plastically tune trait expression to the state of the resource budget or the pace of resource acquisition, releasing maturation from developmental constraints and thereby accommodating optimal maturation timing. This hypothesis, which we coin ‘plastic release hypothesis’, or ‘polyphenic release hypothesis’ in cases of a sexual polyphenism, remains untested, despite its potential for understanding the poorly explored relationship between sexual traits and life history^26^. A deeper understanding of how sexual trait development impacts the timing of life history events, and vice versa, could be an important step towards unravelling the contribution of developmental and physiological constraints to life history evolution^3,6,9,27^.

Here, we test the polyphenic release hypothesis in bulb mites (*Rhizoglyphus robini*). These mites form large populations on ephemeral food sources such as rotting plant roots, and experience highly variable levels of competition for food as they go through boom-bust cycles^28^. Hence, age at maturity is strongly plastic and nutrition-dependent, ranging from less than a week in well-fed large individuals to more than a month in poorly-fed small individuals^29^. Additionally, *Rhizoglyphus* mites show a conspicuous polyphenism in a male secondary sexual trait^30^. Large males typically mature as a ‘fighter’, armed with dagger-like tarsal claws at the end of a greatly enlarged third leg pair (Fig. 1a) that they use to puncture the cuticle of conspecifics, thereby killing them. Smaller males mature as an unarmed ‘scrambler’ with unmodified legs adapted for walking (Fig. 1b)^31^. In *R. robini*, scrambler expression is associated with poor growth immediately before metamorphosis (Rhebergen *et al.* in prep). In several related acarid mites that share the fighter-scrambler polyphenism, scrambler expression is additionally induced by pheromones from dense colonies^32,33^; this is not the case in *R. robini*^32^.

**Figure 1.**
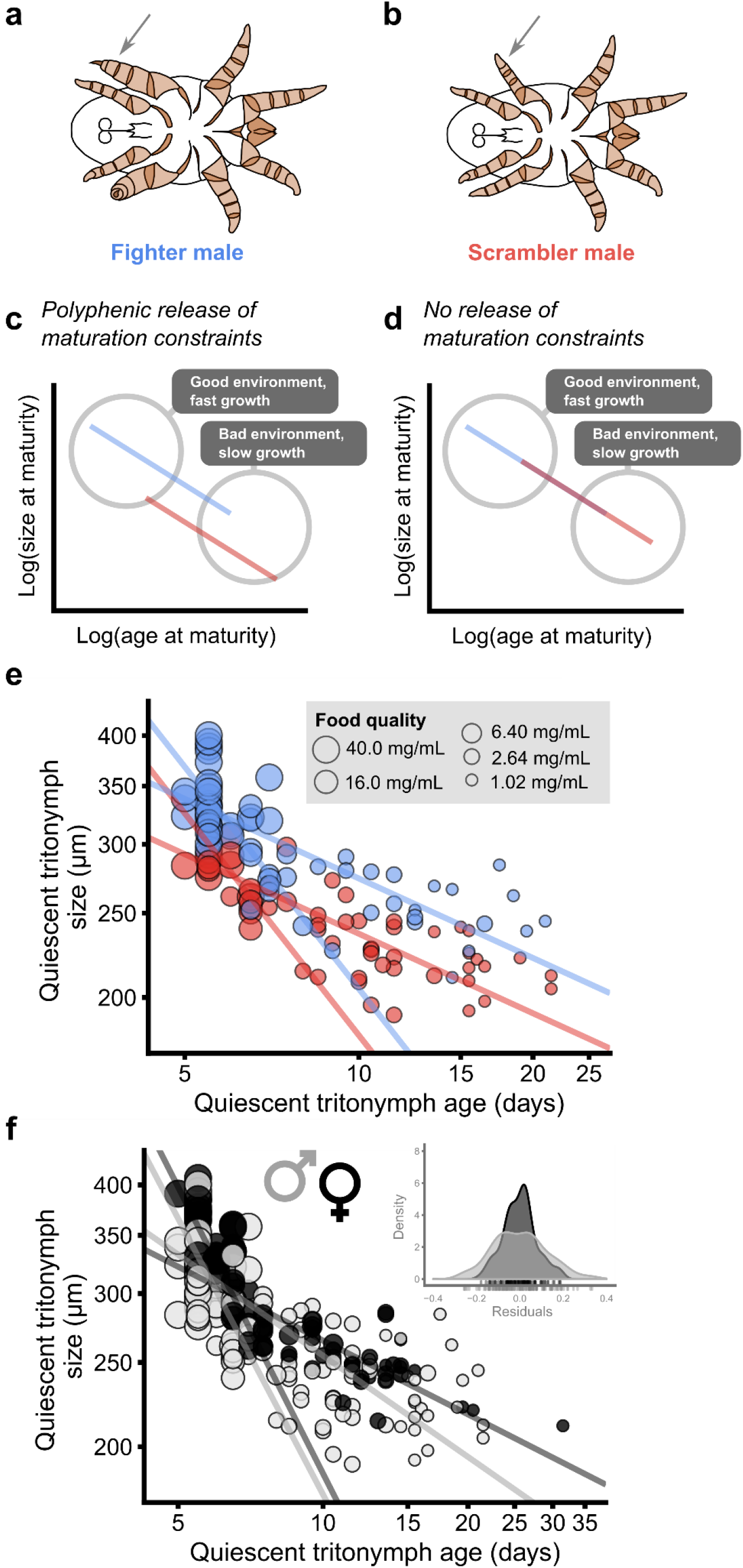
Reaction norms of age and size at maturation for *Rhizoglyphus robini* male morphs. **a,** The male fighter morph. The third leg pair is strongly modified and enlarged (arrow). **b,** The male scrambler morph. The third leg pair is not modified (arrow). **c,** The polyphenic release hypothesis predicts that the reaction norm of age and size at maturity is elevated in fighters (blue), relative to scramblers (red). Axes are log-transformed to linearize the generally strongly L-shaped relationship between age and size at maturity^3,29^. **d,** Alternatively, if fighter expression does not constrain maturation timing, age and size at maturity will fall on the same curve for fighters and scramblers. **e,** Age and size at metamorphosis (log-transformed) of fighters (blue) and scramblers (red) under variation in food quality (circle width). The fighter reaction norm is elevated compared to the scrambler reaction norm (fast-growing males: Wald 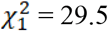, P < 0.001; slow-growing males: Wald 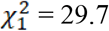, P < 0.001). Lines represent SMA regressions for fast- and slow-growing fighters and scramblers separately. **f,** Age and size at metamorphosis (log-transformed) of males (grey) and females (black) under variation in food quality (circle width). Males follow the average norm of reaction less closely than females: the distribution of SMA residuals is wider for males than for females (F_126,109_ = 2.67, P < 0.001). Lines represent SMA regressions for fast- and slow-growing males and females separately.

According to the polyphenic release hypothesis, if enlarged fighter legs are physiologically expensive to produce, the fighter-scrambler polyphenism lowers the physiological threshold for entering metamorphosis and maturation by facultatively removing a major resource allocation target (weaponry). Here, combining individual-level growth manipulations with population-level manipulations of the strength of competition for food, we investigate whether scrambler expression can prevent juvenile male *R. robini* from being trapped in the juvenile stage when resource acquisition is slow.

## Results

In our first experiment, we tested the prediction of the polyphenic release hypothesis that scrambler expression enables males to enter metamorphosis at an earlier age (and hence smaller size) than fighters on the same growth trajectory. Thus, the reaction norm of age and size at metamorphosis in response to variation in somatic growth rate should be elevated in fighters relative to scramblers; the difference in elevation being evidence for a sexually selected constraint imposed on the reaction norm (Fig. 1c,d). We also tested the assumption that fighter expression is physiologically more expensive than scrambler expression, particularly for poorly-grown males. We generated variation in growth rates by rearing males to adulthood on disks of filter paper soaked in a yeast concentration of 40.0 mg/mL (n = 35), 16.0 mg/mL (n = 21), 6.40 mg/mL (n = 12), 2.56 mg/mL (n = 37) and 1.02 mg/mL (n = 23). We recorded age and size at metamorphosis (i.e. the quiescent tritonymph stage), size change after metamorphosis (as a proxy for physiological cost of adult phenotype development that tests the assumption of fighter expression being costly), and male morph.

Fighters (n = 64) shrank on average significantly more than scramblers (n = 60) during metamorphosis (likelihood ratio test (LRT) 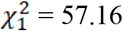, P < 0.001); the average fighter lost 6.1% of its body size (95% CI [5.1, 7.1]; Fig. 2), whereas the average scrambler lost only 2.0% of its body size (95% CI [1.0, 3.0]; Fig. 2). Shrinking was less for larger quiescent tritonymphs (LRT 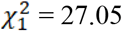, P < 0.001); average shrinkage decreased 3.4 percentage points with each 100 μm body width (95% CI [2.2, 4.6]; Fig. 2). Consequentially, the largest fighters did not shrink, whereas the smallest fighters lost up to 11.8% of their body size at the transition from juvenile to adult, supporting our assumption that the fighter morph is costly to develop for small males, and subject to resource allocation trade-offs. Such physiological costs may translate to substantial fitness costs, if only because body size could be an important determinant of mating success in fighters^34^. The interaction between quiescent tritonymph body size and male morph did not significantly affect shrinking (LRT 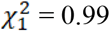, P = 0.320).

**Figure 2.**
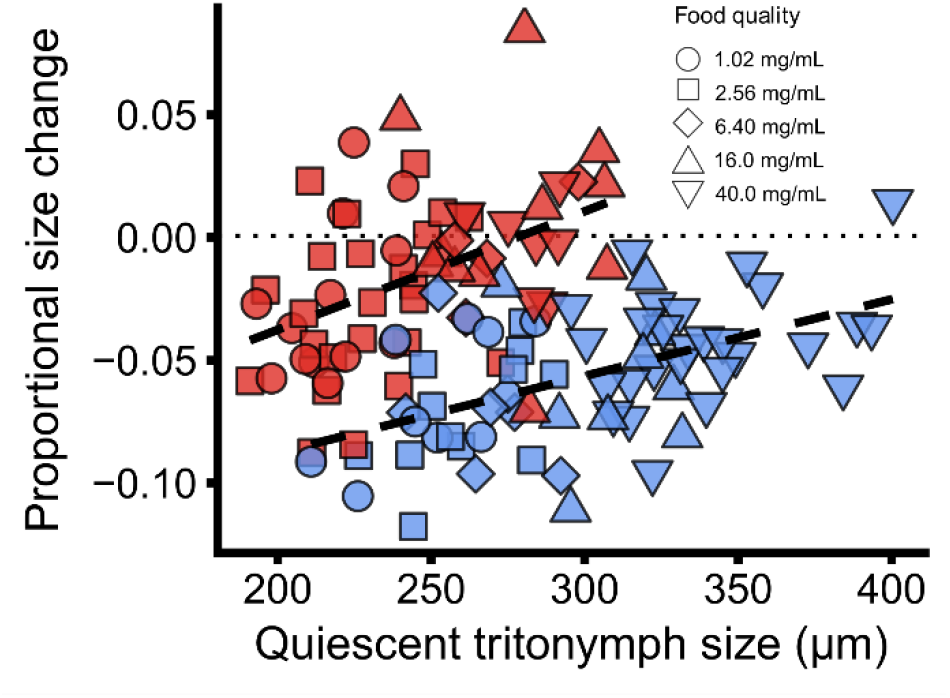
Proportional shrinking or growing during male metamorphosis, as a function of body size at metamorphosis and male morph. Variation in quiescent tritonymph body size (i.e. size at metamorphosis) was experimentally induced by varying food quality (symbol shapes). Dashed lines represent the regression lines for fighters and scramblers. Four intermorph males are not shown in this figure. Proportional size change depended on size at metamorphosis (linear mixed model, LRT 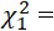 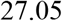, P < 0.001), and fighters shrank more than scramblers (LRT 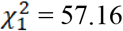, P < 0.001).

The reaction norm of age and size at maturity did not significantly differ in slope for fighter and scrambler males (fast-growing males: LRT 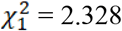, P = 0.127; slow-growing males: LRT 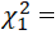 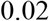 = 0.02, P = 0.877), but, in line with our prediction, was consistently shifted upwards for fighter males, compared to scramblers (fast-growing males: Wald 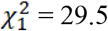, P < 0.001; slow-growing males: Wald 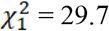, P < 0.001; Fig. 1e). Fast-growing males (40 μm/day, on the fast end of the spectrum) accelerated maturation by 0.4 days, and slow-growing males (8 μm/day, on the slow end of the spectrum) by 2.7 days, if they matured as scramblers rather than fighters. For slow-growing males in adverse circumstances when juvenile mortality is high, this represents a considerable timing advantage, presumably due to polyphenic release of developmental constraints. As a consequence, the residual distribution of body sizes at metamorphosis was wider for males than for females, which, unlike males, cannot switch to an energetically cheaper adult phenotype during metamorphosis and thus follow the average reaction norm more closely (F_126,109_ = 2.67, P < 0.001; Fig. 1f).

In interpreting the results from our first experiment, we assumed that bulb mites adaptively time the onset of metamorphosis cued by current, local circumstances, such as a perceived worsening of growth conditions^4,5,35^. Under this assumption, polyphenic release from sexually selected constraints could be a powerful developmental strategy allowing early metamorphosis with even a very small energy budget. Thus, in our second experiment, we tested the hypothesis that the onset of metamorphosis is sensitive to diminishing food conditions. We predicted this effect to be stronger in males than in females, because males can plastically switch to a cheaper phenotype during metamorphosis, whereas females cannot. We reared freshly emerged tritonymphs (the final feeding juvenile stage) to adulthood on *ad lib* access to yeast, removed their food after 26 hours (n=23 males, 27 females), 30 hours (n=20 males, 29 females), 34 hours (n=14 males, 32 females) or not at all (n=22 males, 28 females) and recorded metamorphosis timing, sex, and adult male morph. We found that both males and females accelerated metamorphosis onset in response to food removal (Fig. 3a-d). After 36 hours, only 14% of the mites had entered metamorphosis in the non-removal treatment, whereas 30%, 53% and 44% had entered metamorphosis in the 34 hours, 30 hours and 26 hours treatment, respectively (LRT 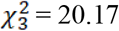, P < 0.001; Fig. 3b,d). The sexes did not significantly differ in their response to food removal (LRT 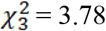, P = 0.286; Fig. 3b,d), suggesting that females may employ developmental mechanisms of their own to facilitate accelerated metamorphosis. The sexes did not significantly differ in average metamorphosis timing (LRT 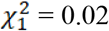, P = 0.878). We did not detect an effect of food removal on male morph expression (LRT 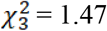, P = 0.689).

**Figure 3.**
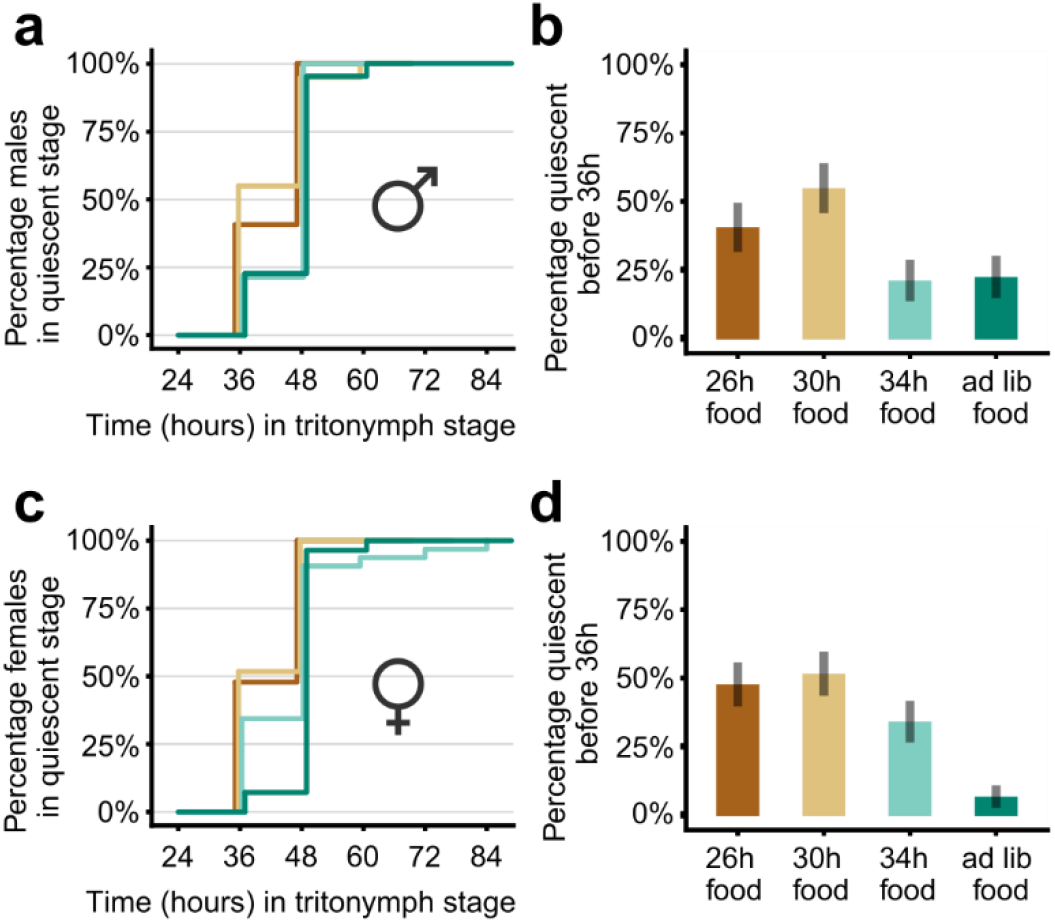
Mites accelerate maturation in response to food removal. Percentage of male **(a)** and female **(c)** tritonymphs in metamorphosis over time, in four treatments: food removed after 26 hours in the tritonymph stage (dark brown), 30 hours (light brown), 34 hours (light green) or not at all (dark green). Percentage of male **(b)** and female **(d)** tritonymphs that had entered metamorphosis after less than 36 hours in the tritonymph stage, in response to food removal treatment. Error bars denote standard errors of the proportion. The percentage of tritonymphs in metamorphosis after less than 36 hours depended significantly on food removal regime (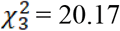, P < 0.001) but not on sex (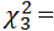 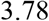, P = 0.286).

If polyphenic release is an adaptation to variable and occasionally unfavourable environments, it should result in an adaptive response to adverse population conditions. We tested this prediction in our third experiment. If competition for food is experimentally increased in laboratory populations, increasingly more males should express the scrambler morph rather than the fighter morph, because scramblers can mature earlier and at a smaller body size than fighters on the same growth trajectory. We prepared populations of newly hatched larvae in four treatments: low population density (15 larvae) and low food availability (two yeast rods, 0.25 mg; hereafter ‘LL treatment’), high population density (40 larvae) and high food availability (10 yeast rods, 1.25 mg; hereafter ‘HH treatment’), high population density and low food availability (hereafter ‘HL treatment’), and individually isolated larvae (40 larvae) with *ad lib* yeast (hereafter ‘isolated treatment’). Hence, competition for food was high in the HL treatment, and low in the other three treatments. We repeated this setup three times. We recorded size and age at the onset of metamorphosis (the quiescent phase) in each juvenile stage (larvae, protonymphs, tritonymphs), and we recorded sex and male morph in surviving adults. The effect of treatment on mortality changed with experimental replicate (LRT 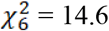, P = 0.024). Over all replicates, however, mortality was considerably higher in the LH treatment (38%, Fig. 4a) than in the LL, HH and isolated treatments (9%, 4% and 4% respectively; Fig. 4a). Likewise, size at metamorphosis depended on treatment in all three juvenile stages (larvae: Kruskal-Wallis 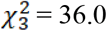, P < 0.001; protonymphs: Kruskal-Wallis 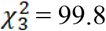, P < 0.001; male tritonymphs: Kruskal-Wallis 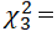 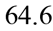, P < 0.001; female tritonymphs: Kruskal-Wallis 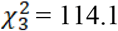, P < 0.001; Fig. 4b), which was mainly due to poor growth in the HL treatment, but also the increased growth of females in the isolated treatment. Scrambler expression strongly responded to population treatment (LRT 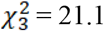, P < 0.001), and was higher in the HL treatment (77%; Fig. 4c) than in the LL, HH and isolated treatments (32%, 41% and 28% respectively; Fig. 4c). This effect did not depend on experimental replicate (LRT 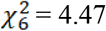, P = 0.614). The difference between fighters and scramblers in the relationship between age and size at metamorphosis in the population experiment, pooled across all treatments, mirrored the difference in the reaction norms observed in our first experiment. The slope was not significantly different for fighters and scramblers (LRT 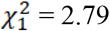, P = 0.095) but fighters consistently matured later and at a larger size than scramblers on the same growth trajectory (Wald 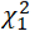 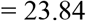, P < 0.001; Fig. 5). Bilaterally asymmetric ‘intermorph’ males, which were scrambler on one side and fighter on the other, occurred only in the HH treatment (n=5 over all replicates).

**Figure 4.**
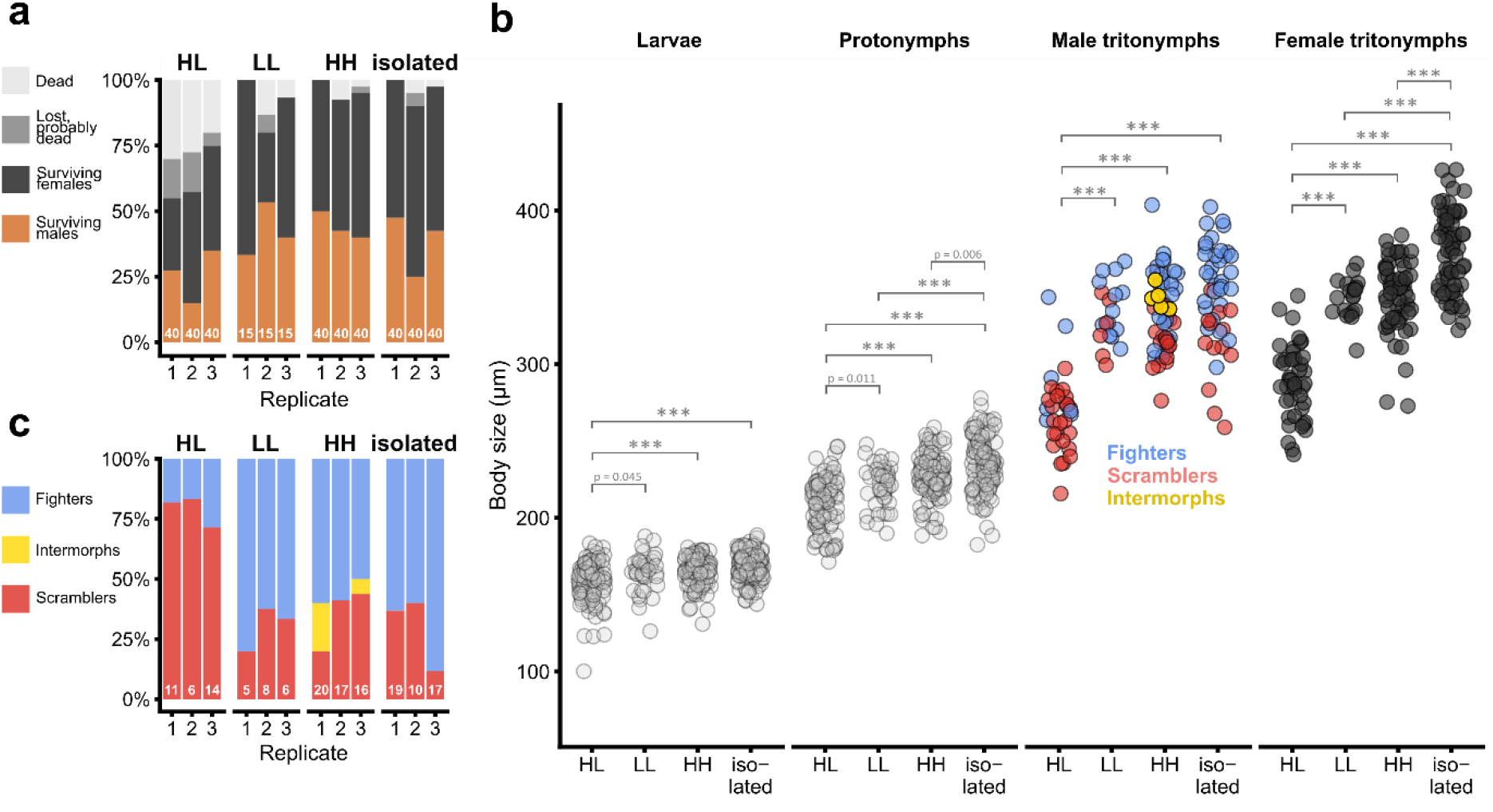
Mortality, growth and male morph expression under competition for food. **a,** Mortality in four competition treatments: high population density, low food availability (HL); low population density, low food availability (LL); high population density, high food availability (HH); and individually isolated with *ad libitum* food (isolated). Numbers at the bottom of each bar denote initial population sizes. **b,** Body size (maximal idiosoma width) of quiescent larvae, protonymphs and tritonymphs in the four competition treatments, all replicates pooled. Size at quiescence differed among treatments (larvae: Kruskal-Wallis 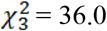, P < 0.001; protonymphs: Kruskal-Wallis 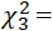 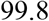, P < 0.001; male tritonymphs: Kruskal-Wallis 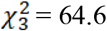, P < 0.001; female tritonymphs: Kruskal-Wallis 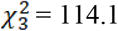, P < 0.001). Brackets denote statistically significant post-hoc pairwise comparisons (Nemenyi’s post-hoc test); all *p < 0.001* except where indicated. **c,** Male morph expression depended on competition treatment (LRT 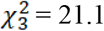, P < 0.001). Numbers at the bottom of each bar denote the number of males that had survived until adulthood in each replicate.

**Figure 5.**
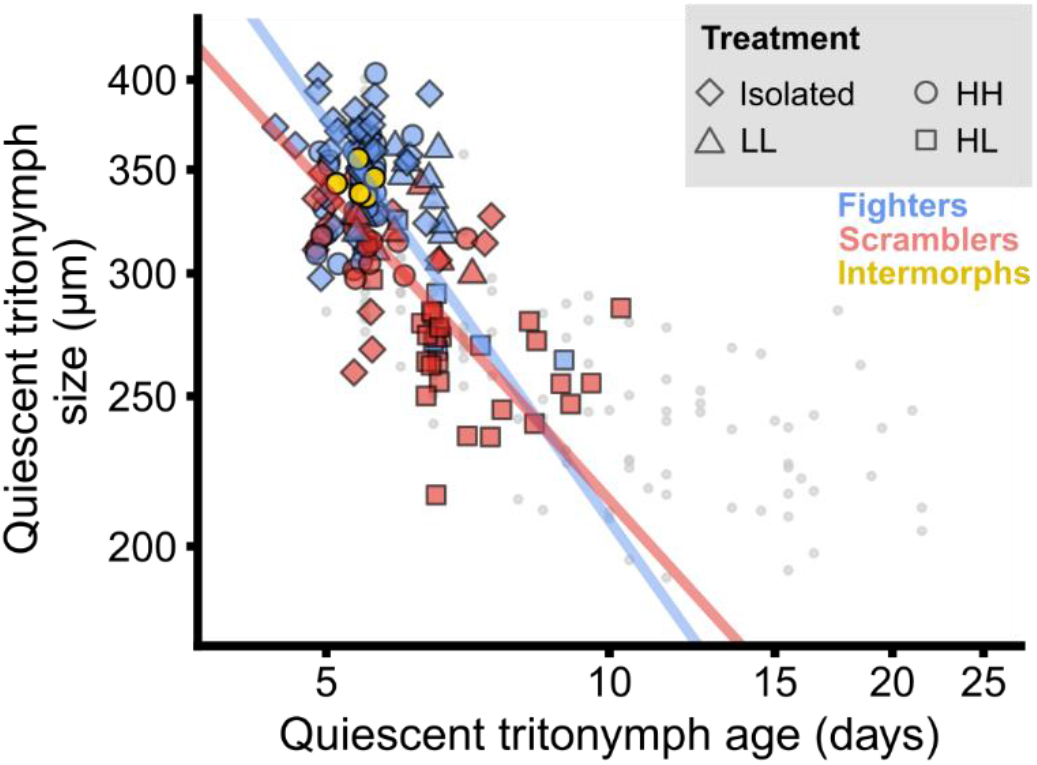
Observed relationship between age and size at metamorphosis (log-transformed) in fighters and scramblers, under variation in competition for food. The SMA regression line for fighters is elevated compared to the regression line for scramblers (Wald 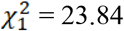, P < 0.001). Different shapes denote competition treatments: rectangle, high population density, low food availability (HL); triangle, low population density, low food availability (LL); circle, high population density, high food availability (HH); and diamond, individually isolated with *ad libitum* food (isolated). Light grey points show the male reaction norm of age and size at metamorphosis as observed in experiment 1 (Fig. 1e).

## Discussion

Our results provide the first evidence for polyphenic release of developmental maturation constraints. Slow-growing males were quicker to escape the risky juvenile stage, and avoided a considerable amount of shrinking (in line with earlier findings^36^), if they did not develop a morphologically exaggerated trait. Based on this result, we hypothesize that scrambler expression is a consequence of adaptive re-allocation of resources to vital processes during metamorphosis^37–39^, which is necessary when adverse environmental circumstances favour early maturation with a small resource budget^4,5,40^. Thus, it may not be the scrambler morph *itself* that is adaptive, but rather the resource allocation switch underlying its condition-dependent expression. An alternative but equally intriguing explanation for our results is that poorly growing males anticipate poor future fighting success, refrain from developing into a fighter, and accelerate maturation for an adaptive sneaking tactic^30^. Accordingly, scrambler expression and accelerated maturation could both be part of an adaptive sneaker syndrome. There is currently no evidence that scramblers employ distinct sneaking behaviour to gain access to females, apart from their inability to kill rivals^31^. Furthermore, in small males, the scrambler morph does not appear to improve fertilization success compared to the fighter morph^41^. This renders the ‘adaptive sneaker’ hypothesis as yet unsupported, although it is not mutually exclusive with the polyphenic release hypothesis. All in all, scrambler males may truly have to ‘make the best of a bad lot’^42^. They cannot kill, and hence potentially miss out on mating opportunities. However, males that develop as scramblers are quicker to mature. In adverse environments, such accelerated maturation prevents juveniles from dying before reproducing^4,5,40^. This fitness benefit, accrued during early life, may partly, or possibly completely, compensate the fitness costs of not carrying weaponry later in the life cycle. The resulting life history flexibility may have contributed to the cosmopolitan spread of *Rhizoglyphus* mites, as well as their status as a major agricultural pest^28^.

The selection pressures determining optimal maturation timing^1,3,40,43,44^ are often studied independently from the selection pressures that shape sexual trait expression^45,46^. Yet, our results imply that the evolutionary dynamics of secondary sexual traits and life history traits are likely intertwined^26,47^. Trait exaggeration due to sexual selection inevitably causes trait development to depend on an increasingly large portion of the energy budget^26,48,49^, and timing of life history events is typically energy-budget-dependent^50^. Indeed, comparative evidence in birds^51^ and fruit flies^52–54^ suggests that the evolution of exaggerated secondary sexual trait results in prolonged maturation time, presumably because development of such exaggerated traits is energetically expensive. Hence, we need to consider the possibility that flexibility in the timing of life history events evolves by introducing plasticity in energetically expensive morphological traits, or that pre-existing plasticity in morphology facilitates the evolution of life history flexibility. To our knowledge, our results provide the first evidence that facultative expression of a male trait promotes flexible maturation timing through polyphenic release of developmental constraints.

## Methods

### Study system and general procedures

The bulb mite *Rhizoglyphus robini* (Acari: Acaridae) is a common cosmopolitan mite species, which naturally forms dense colonies on rotting plant roots and flower bulbs^28^. The populations used in this study are derived from *R. robini* samples that were collected from flower bulbs in horticultural storage rooms in Noord-Holland, the Netherlands, in December 2010. *R. robini* normally has three juvenile stages (larva, protonymph, tritonymph), interspersed with quiescent phases during which individuals do not move or eat but undergo incomplete metamorphosis to enter the next stage^28^. Maturation occurs immediately after the quiescent tritonymph stage (in this study referred to as: metamorphosis). In extremely adverse conditions, juveniles go through a rare fourth stage called the deutonymph, which is adapted for phoretic dispersal and is expressed between the protonymph and tritonymph stage^28^. None of the mites in our study expressed the deutonymph stage.

We keep our *R. robini* populations in an unlit climate cabinet (25 °C; >90% humidity), in plastic containers (10 x 10 x 2.5 cm) with a plaster of Paris substrate, on a diet of baker’s yeast. Under these conditions, generation time averages 11 days. Twice a week, we clean one sixth of the substrate by removing all organic matter and add fresh water and food. We mix our populations every six months to maintain genetic homogeneity.

In the experiments, individual mites were kept in individual plastic tubes (height 5 cm, diameter 1.5 cm) with a moistened plaster substrate. The tubes were sealed with fine mesh, kept in place by a plastic lid and placed in an unlit climate cabinet (25 °C; >90% humidity) throughout the experiment. Experimental groups of mites (experiment 3) were kept in wider tubes (height 5 cm, diameter 2.5 cm). Mites were photographed ventrally and measured to the nearest 0.001 μm using a Zeiss Axiocam 105 color camera mounted on a Zeiss Stemi 200-C stereomicroscope, and ZEN 2 (Blue edition) software.

For statistical analysis and graphs, we used R v. 3.6.1^55^, including the packages *stats, ggplot2*^56^*, dplyr*^57^*, smatr*^58^*, segmented*^59^ and *PMCMR*^60^.

### Experiment 1

The results of experiment 1 form part of an experiment of which the methods are reported elsewhere (Rhebergen *et al.* in prep.). For clarity, we briefly report the methods here as well. The parental generation was obtained by individually isolating 200 larvae from the yeast-fed stock populations, in four experimental blocks of 50 larvae each. These larvae were reared to adulthood on a diet of *ad libitum* yeast in individual tubes, to eliminate bias due to parental effects. Per block, 15-24 virgin adult males and females of the parental generation were randomly paired in a plastic tube with *ad libitum* yeast to produce the focal generation. Three days after pairing, eggs of each parental pair were transferred to a clean individual tube without food. Within 12 hours after hatching, larvae were transferred to experimental tubes. Each parental pair contributed a single larva to each food treatment. Note that, to account for genetic variation associated with life history traits and male morph expression, we sampled the parental generation randomly from our stocks. An isoline approach to test several genotypes explicitly would have been logistically infeasible.

The focal individuals were individually isolated and reared to adulthood on quartered ~23 mm^2^ circular disks of filter paper that were soaked for 20 seconds in 10 μL solution of yeast in water. Individuals were subjected to five food treatments, which represented a 2.5-fold dilution series (yeast solved in water): 40.0 mg/mL (n = 69), 16.0 mg/mL (n = 40), 6.40 mg/mL (n = 40), 2.56 mg/mL (n = 69) and 1.02 mg/mL (n = 40). Experimental blocks 1 and 4 contained all treatments, but due to logistical constraints, blocks 2 and 3 only contained the 40.0 mg/mL and 2.56 mg/mL treatments. Every two days, the substrate of each tube was cleaned with a moist brush, and the yeast-soaked disk was replaced with a fresh disk to prevent mould from growing.

We checked whether individuals had reached the quiescent tritonymph stage every 12 hours. Once individuals had done so, we scored age at metamorphosis (in 12h intervals) and photographed each individual ventrally. To score size at metamorphosis, we measured the width of the idiosoma (the white bulbous posterior part of the body; measurement st. dev. = 3.3 μm; Supplementary Information). Roughly 24 hours later, when adults had emerged, we recorded sex and male morph, and photographed each individual again to measure size after metamorphosis. Of the 258 focal individuals, 17 died as a juvenile, yielding a total of 128 adult males and 113 adult females. We failed to obtain a quiescent tritonymph measurement of 1 scrambler male and 3 females.

To test whether (proportional) size change during metamorphosis depends on male morph and body size at metamorphosis, we constructed a linear mixed-effects model with proportional body size change during metamorphosis as response variable. Fixed explanatory variables were idiosoma width at metamorphosis, male morph (fighter or scrambler; 4 intermorph males were excluded from the analysis), the interaction between idiosoma width and male morph, and block. Parent identity was included as random intercept. We defined the response variable, proportional change in idiosoma width, as *[adult idiosoma width –idiosoma width at metamorphosis] / idiosoma width at metamorphosis*. We centred the explanatory variable ‘idiosoma width at metamorphosis’ around the mean value for fighters and scramblers separately, so that the intercept and morph parameters directly corresponded to the average amount of proportional shrinking in fighters and scramblers. We tested the assumptions of linearity, normality and homoscedascity by inspecting plots of the residuals against the fitted values and the explanatory variables, and by inspecting a normal quantile-quantile plot of the residuals. We tested significance of model parameters using likelihood ratio tests (LRT).

To assess whether the reaction norm of age and size at metamorphosis differed in slope and elevation between fighters and scramblers, we performed standardized major axis (SMA) regression on the log-transformed age and size at metamorphosis of males (comprising fighters and scramblers; 4 intermorph males were excluded from this analysis), using the *smatr* package in R, with male morph as a grouping factor. As preliminary analyses indicated that the reaction norm was non-linear even when log-transformed, we first performed a breakpoint analysis using the *segmented* package to find the metamorphosis age at which the reaction norm changed. We then compared the breakpoint linear model with a regular linear model using the Akaike Information Criterion (AIC), and inspected the residuals of the breakpoint model visually to exclude remaining non-linearity. The breakpoint model performed considerably better than the linear model (∆AIC = 20.2), so we proceeded by fitting SMA regressions for fast-growing and slow-growing males separately. Following Warton *et al.*^58^, we tested for a difference in the slope of the fighter and scrambler reaction norms using a likelihood ratio test (LRT), and for a difference in elevation using a Wald test. Finally, we compared the degree to which age and size at metamorphosis followed the average reaction norm in males (including the four intermorphs) and females, by fitting SMA regressions for slow- and fast-growing males and females separately, extracting the residuals, and comparing the variance of the male and female residuals using an F-test.

### Experiment 2

We randomly collected 200 protonymphs from our stock populations over two days, and isolated them in individual tubes with *ad libitum* yeast. Once they had entered the quiescent protonymph stage, we removed their food. Every morning, we checked for freshly emerged tritonymphs, which emerged over the course of three days in tubes without food. The freshly emerged tritonymphs were randomly allocated to one of four treatments: 1) *ad libitum* access to food (yeast), but food removed after 26 hours (n=50); 2) *ad libitum* access to food, but food removed after 30 hours (n=50); 3) *ad libitum* access to food, but food removed after 34 hours (n=50); and 4) *ad libitum* access to food until metamorphosis (n=50). Every 12 hours, we scored whether tritonymphs had entered metamorphosis. Roughly 24 hours after the onset of metamorphosis, when adults had emerged, we scored sex and adult male morph.

The tritonymphs were observed to have entered metamorphosis on a limited number of time points, and the vast majority on only two time points (36h and 48h after the start of the treatments). Therefore, we rescored the time of metamorphosis as ‘early metamorphosis’ (36h since the start of the treatment) or ‘late metamorphosis’ (>36h since the start of the treatment). To assess whether treatment influenced the proportion of tritonymphs entering metamorphosis early, we performed a logistic regression (generalized linear model) with metamorphosis timing (early or late) as binary response variable, and treatment, sex and day of tritonymph emergence as fixed explanatory variables. We used LRT to assess whether treatment, sex and their interaction had statistically significant effects.

### Experiment 3

The population experiment was done in three replicates, each replicate containing each of the four treatments once. For each replicate, the parental generation was obtained following the same procedure as in experiment 1, except we collected 150 larvae from the stock populations per experimental replicate, forming 50 virgin male-female pairs to produce the focal generation. We isolated eggs following the procedure of experiment 1. Within 12h after hatching, a single larva per parental pair was transferred to each experimental treatment, to maintain genetic homogeneity across the treatments within each replicate.

Per experimental replicate, we prepared three population tubes containing three population treatments (HH: high population density, high food availability; HL: high population density, low food availability, and LL: low population density, low food availability). Additionally, we prepared 40 individual tubes for our fourth experimental treatment (individually isolated, no population effects). In the HH tube, we added 1.25 mg baker’s yeast, which corresponded to 10 yeast rods. In the HL, LL and isolated tubes, we added 0.25 mg baker’s yeast, which corresponded to 2 yeast rods. The HH, HL and isolated treatments received 40 newly hatched larvae each, which were placed on top of the food source using a fine moist brush. The LL treatment received 15 larvae. To prevent mould from growing, the population treatments (HH, HL and LL) were transferred to fresh tubes first after 3 days, then every 2 days, by carefully picking up each individual mite with a moist brush and putting them in a freshly prepared tube with fresh food in the quantities described above. Hence, the HH, HL and LL treatments received a pulse of fresh food first after 3 days, then every 2 days. The mites in the isolated treatment were not transferred to fresh tubes, but did receive fresh food at the same time as the population tubes.

We monitored all the tubes every 12 hours, and recorded the number of larvae, protonymphs and tritonymphs, the number of dead individuals and the number of individuals that had entered the quiescent phase. We photographed all quiescent larvae, protonymphs and tritonymphs ventrally and measured the maximal width of the idiosoma (the large bulbous posterior part of the body) as a proxy for body size. We collected and isolated all quiescent tritonymphs in individual tubes, and recorded sex and male morph when the adult emerged. We then put the adult back in the population tube where it came from.

To assess survival until maturity, we performed a logistic regression (generalized linear model) with survival (yes/no) as binary response variable (we assumed lost individuals to be dead), and treatment and replicate as fixed explanatory variables. To assess the effect of treatment on male morph expression, we performed a logistic regression with male morph as binary response variable (note that 5 ‘intermorph’ males were omitted from this analysis), and treatment and replicate as fixed explanatory variables. We used LRT to assess whether survival and male morph expression were significantly affected by treatment, and whether the effect changed with experimental replicate. To assess the effect of treatment on quiescent larval, protonymphal and tritonymphal body size, we used the non-parametric Kruskal-Wallis analysis of variance, as preliminary analyses showed evidence for non-normality and heteroscedascity. We used the Nemenyi post-hoc test for pairwise comparison of treatments.

Finally, to assess whether the reaction norm of age and size at metamorphosis differed in slope and elevation between fighters and scramblers, we performed standardized major axis (SMA) regression on the log-transformed age and size of male quiescent tritonymphs (pooled from all treatments and replicates), with male morph as a grouping factor. We tested for a difference in the slope of the fighter and scrambler reaction norms using a likelihood ratio test (LRT), and for a difference in elevation using a Wald test, following Warton *et al.*^58^.

## Supporting information

Analysis of measurement error

## Acknowledgements

This work is funded by a VIDI grant (No. 864.13.005) from the Netherlands Organisation for Scientific Research (NWO) awarded to IMS. Fabian Sterken did an initial pilot study as part of his BSc degree, which prompted this work.

## References

1. Stearns, S. C. & Koella, J. C. The Evolution of Phenotypic Plasticity in Life-History Traits: Predictions of Reaction Norms for Age and Size at Maturity. Evolution 40, 893 (1986).

2. Teder, T., Vellau, H. & Tammaru, T. Age and size at maturity: A quantitative review of diet-induced reaction norms in insects. Evolution 68, 3217–3228 (2014).

3. Day, T. & Rowe, L. Developmental thresholds and the evolution of reaction norms for age and size at life-history transitions. Am. Nat. 159, 338–350 (2002).

4. Wilbur, H. M. & Collins, J. P. Ecological aspects of amphibian metamorphosis. Science 182, 1305–1314 (1973).

5. Morey, S. & Reznick, D. A Comparative Analysis of Plasticity in Larval Development in Three Species of Spadefoot Toads. Ecology 81, 1736 (2000).

6. Nilsson-Örtman, V. & Rowe, L. The evolution of developmental thresholds and reaction norms for age and size at maturity. Proc. Natl. Acad. Sci. U. S. A. 118, (2021).

7. Blanckenhorn, W. U. The Evolution of Body Size: What Keeps Organisms Small? The Quarterly Review of Biology 75, 385–407 (2000).

8. Berner, D. & Blanckenhorn, W. U. An ontogenetic perspective on the relationship between age and size at maturity. Funct. Ecol. 21, 505–512 (2007).

9. Ricklefs, R. & Wikelski, M. The physiology/life-history nexus. Trends Ecol. Evol. 17, 462–468 (2002).

10. West-Eberhard, M. J. Developmental Plasticity and Evolution. (Oxford University Press, 2003).

11. Moran, N. A. Adaptation and constraint in the complex life cycles of animals. Annu. Rev. Ecol. Syst. 25, 573–600 (1994).

12. Ten Brink, H., de Roos, A. M. & Dieckmann, U. The evolutionary ecology of metamorphosis. Am. Nat. 193, E116–E131 (2019).

13. Rolff, J., Johnston, P. R. & Reynolds, S. Complete metamorphosis of insects. Philos. Trans. R. Soc. B Biol. Sci. 374, 20190063 (2019).

14. Nijhout, H. F. & Emlen, D. J. Competition among body parts in the development and evolution of insect morphology. Proc. Natl. Acad. Sci. 95, 3685–3689 (1998).

15. Emlen, D. J. The Evolution of Animal Weapons. Annu. Rev. Ecol. Evol. Syst. 39, 387–413 (2008).

16. Wiens, J. J. & Tuschhoff, E. Songs versus colours versus horns: what explains the diversity of sexually selected traits? Biol. Rev. 7, (2020).

17. Zahavi, A. Mate selection-a selection for a handicap. J. Theor. Biol. 53, 205–14 (1975).

18. Getty, T. Sexually selected signals are not similar to sports handicaps. Trends Ecol. Evol. 21, 83–8 (2006).

19. Emlen, D. J. Costs and the diversification of exaggerated animal structures. Science 291, 1534–6 (2001).

20. O’Brien, D. M. et al. Muscle mass drives cost in sexually selected arthropod weapons. Proc. R. Soc. B Biol. Sci. 286, (2019).

21. Warren, I. A., Gotoh, H., Dworkin, I. M., Emlen, D. J. & Lavine, L. C. A general mechanism for conditional expression of exaggerated sexually-selected traits. BioEssays 35, 889–899 (2013).

22. Tomkins, J. L. Environmental and genetic determinants of the male forceps length dimorphism in the European earwig Forficula auricularia L. Behav. Ecol. Sociobiol. 47, 1–8 (1999).

23. Smallegange, I. M. Complex environmental effects on the expression of alternative reproductive phenotypes in the bulb mite. Evol. Ecol. 25, 857–873 (2011).

24. Moczek, A. P. Horn polyphenism in the beetle Onthophagus taurus: larval diet quality and plasticity in parental investment determine adult body size and male horn morphology. Behav. Ecol. 9, 636–641 (1998).

25. Cotton, S., Fowler, K. & Pomiankowski, A. Do sexual ornaments demonstrate heightened condition-dependent expression as predicted by the handicap hypothesis? Proc. R. Soc. B 271, 771–83 (2004).

26. Morehouse, N. I. Condition-dependent ornaments, life histories, and the evolving architecture of resource-use. Integr. Comp. Biol. 54, 591–600 (2014).

27. Zera, A. J. & Harshman, L. G. The Physiology of Life History Trade-Offs in Animals. Annu. Rev. Ecol. Syst. 32, 95–126 (2001).

28. Díaz, A., Okabe, K., Eckenrode, C. J., Villani, M. G. & Oconnor, B. M. Biology, ecology, and management of the bulb mites of the genus Rhizoglyphus (Acari: Acaridae). Exp. Appl. Acarol. 24, 85–113 (2000).

29. Smallegange, I. M. Effects of paternal phenotype and environmental variability on age and size at maturity in a male dimorphic mite. Naturwissenschaften 98, 339–346 (2011).

30. Radwan, J. W. Alternative mating tactics in acarid mites. in Advances in the Study of Behavior, Vol. 39 185–208 (Elsevier Academic Press, 2009).

31. Radwan, J. W. & Klimas, M. Male dimorphism in the bulb mite, Rhizoglyphus robini: fighters survive better. Ethol. Ecol. Evol. 13, 69–79 (2001).

32. Radwan, J. W. Male morph determination in two species of acarid mites. Heredity 74, 669–673 (1995).

33. Radwan, J. Male morph determination in Rhizoglyphus echinopus (Acaridae). Exp. Appl. Acarol. 25, 143–149 (2001).

34. McCullough, E. L. & Simmons, L. W. Selection on male physical performance during male-male competition and female choice. Behav. Ecol. 27, 1288–1295 (2016).

35. Helm, B. R., Rinehart, J. P., Yocum, G. D., Greenlee, K. J. & Bowsher, J. H. Metamorphosis is induced by food absence rather than a critical weight in the solitary bee, Osmia lignaria. Proc. Natl. Acad. Sci. U. S. A. 114, 10924–10929 (2017).

36. Radwan, J. W., Unrug, J. & Tomkins, J. L. Status-dependence and morphological trade-offs in the expression of a sexually selected character in the mite, Sancassania berlesei. J. Evol. Biol. 15, 744–752 (2002).

37. Lea, A. J., Tung, J., Archie, E. A. & Alberts, S. C. Developmental plasticity. Bridging research in evolution and human health. Evol. Med. Public Heal. 2017, 162–175 (2017).

38. Smallegange, I. M., Rhebergen, F. T. & Stewart, K. A. Cross-level considerations for explaining selection pressures and the maintenance of genetic variation in condition-dependent male morphs. Curr. Opin. Insect Sci. 36, 66–73 (2019).

39. Ng’oma, E., Perinchery, A. M. & King, E. G. How to get the most bang for your buck: The evolution and physiology of nutritiondependent resource allocation strategies. Proc. R. Soc. B Biol. Sci. 284, (2017).

40. Rowe, L. & Ludwig, D. Size and timing of metamorphosis in complex life cycles: time constraints and variation. Ecology 72, 413–427 (1991).

41. Michalczyk, Ł., Dudziak, M., Radwan, J. & Tomkins, J. L. Fitness consequences of threshold trait expression subject to environmental cues. Proc. R. Soc. B Biol. Sci. 285, 20180783 (2018).

42. Eberhard, W. G. Beetle Horn Dimorphism: Making the Best of a Bad Lot. Am. Nat. 119, 420–426 (1982).

43. Roff, D. A. Evolution Of Life Histories - Theory and Analysis. (Chapman & Hall, 1993).

44. Stearns, S. C. Life history evolution: successes, limitations and prospects. Naturwissenschaften 87, 476–486 (2000).

45. Andersson, M. & Simmons, L. W. Sexual selection and mate choice. Trends Ecol. Evol. 21, 296–302 (2006).

46. Shuster, S. M. & Wade, M. J. Mating Systems and Strategies. (Princeton University Press, 2003).

47. Cornwallis, C. K. & Uller, T. Towards an evolutionary ecology of sexual traits. Trends Ecol. Evol. 25, 145–152 (2010).

48. Rowe, L. & Houle, D. The lek paradox and the capture of genetic variance by condition dependent traits. Proc. R. Soc. B 263, 1415–1421 (1996).

49. Tomkins, J. L., Radwan, J., Kotiaho, J. S. & Tregenza, T. Genic capture and resolving the lek paradox. Trends Ecol. Evol. 19, 323–328 (2004).

50. Kooijman, S. A. L. M. Dynamic Energy Budget Theory for Metabolic Organisation. (Cambridge University Press, 2010).

51. Ancona, S., Liker, A., Carmona-Isunza, M. C. & Székely, T. Sex differences in age-to-maturation relate to sexual selection and adult sex ratios in birds. Evol. Lett. 1–10 (2020). doi:10.1002/evl3.156

52. Pitnick, S., Markow, T. A. & Spicer, G. S. Delayed male maturity is a cost of producing large sperm in Drosophila. Proc. Natl. Acad. Sci. U. S. A. 92, 10614–8 (1995).

53. Pitnick, S. & Miller, G. T. Correlated response in reproductive and life history traits to selection on testis length in Drosophila hydei. Heredity 84, 416–426 (2000).

54. Pitnick, S., Spicer, G. S. & Markow, T. A. How long is a giant sperm? Nature 375, 109 (1995).

55. R Core Team. R: A language and environment for statistical computing. (2018).

56. Wickham, H. et al. Package ‘ggplot2’: Create Elegant Data Visualisations Using the Grammar of Graphics. (2019).

57. Wickham, H., Francois, R., Henry, L. & Müller, K. Package ‘dplyr’: a Grammar of Data Manipulation. (2018).

58. Warton, D. I., Duursma, R., Falster, D. S. & Taskinen, S. Package ‘smatr’: (Standardised) Major Axis Estimation and Testing Routines. (2018).

59. Muggeo, V. M. R. Package ‘segmented’: Regression Models with Break-Points / Change-Points Estimation. (2017).

60. Pohlert, T. PMCMR: Calculate Pairwise Multiple Comparisons of Mean Rank Sums. (2015).

